# TPR is required for the nuclear export of mRNAs and lncRNAs from intronless and intron-poor genes

**DOI:** 10.1101/740498

**Authors:** Eliza S. Lee, Eric J. Wolf, Harrison W. Smith, Andrew Emili, Alexander F. Palazzo

## Abstract

While splicing has been shown to enhance nuclear export, it has remained unclear whether mRNAs generated from intronless genes use specific machinery to promote their export. Here we investigate the role of the major nuclear pore basket protein, TPR, in regulating mRNA and lncRNA nuclear export in human cells. By sequencing mRNA from the nucleus and cytosol of control and TPR-depleted cells, we provide evidence that TPR is required for the nuclear export of mRNAs and lncRNAs that are generated from intronless and intron-poor genes, and we validate this with reporter constructs. Moreover, in TPR-depleted cells reporter mRNAs generated from intronless genes accumulate in nuclear speckles and are bound to Nxf1. These observations suggest that TPR acts downstream of Nxf1 recruitment, and may allow mRNAs to leave nuclear speckles and properly dock with the nuclear pore. In summary, our study provides one of the first examples of a factor that is required for the nuclear export of intronless and intron-poor mRNAs and lncRNAs.

## Introduction

In eukaryotes mRNA synthesis and processing occurs in the nucleus, while mRNA translation is restricted to the cytoplasm. This subcellular division ensures that only fully processed mRNAs are translated. To accomplish this, mature mRNAs must be recognized and targeted for nuclear export, while pre-mRNAs that have not yet completed splicing must be retained in the nucleus. In addition, cryptic transcripts and misprocessed mRNAs must be nuclear retained and/or degraded.

One of the key components that promote nuclear mRNA export is the Transcription-Export (TREX) complex, which consists of the THOC sub-complex, the mRNA export adaptor Aly/REF and the RNA helicase UAP56 (1–4). This complex is loaded on mRNA and in turn recruits the heterodimeric Nxf1/Nxt1 nuclear transport receptor (also known as TAP/p15) to the mRNA, which helps to ferry it across the nuclear pore (5–9). In addition, a second complex, TREX2, found at the nuclear pore, has also been implicated in mRNA export, although its exact role remains unclear (10–17).

Previously, it was thought that pre-mRNA splicing was a pre-requisite for efficient nuclear mRNA export in metazoans (18, 19). Indeed splicing has been shown to recruit certain TREX components, such as UAP56, to the messenger ribonucleoprotein (mRNP) complex (2, 20, 21). Nonetheless, TREX components have also been shown to be efficiently loaded onto mRNAs even in the absence of splicing (22–26). TREX may be recruited by RNA polymerase II in a co-transcriptional manner (1), by the nuclear cap-binding complex (27) and by the 3’ cleavage and polyadenylation machinery (28). As such, TREX components are required for the export of non-spliced mRNAs (22, 23, 29). In particular cases where splicing enhances nuclear export, it does so by overriding the activity nuclear retention signals (25), although the exact mechanism for how it does so remains unclear. In addition, it has been reported that additional factors are required for the export of naturally intronless mRNAs (30, 31), but to date the role of these factors has not been investigated on a transcriptome-wide scale. Other mRNA features, such as mRNA length may also influence how an mRNA is exported from the nucleus. For example, very short mRNAs (less than 200 nucleotides) do not efficiently recruit the TREX-component Aly and instead are exported by alternative pathways (22, 32).

TPR, one of the main components of the nuclear pore basket (33), helps to tether the TREX2 complex to the nucleoplasmic side of the pore (11, 14, 34). Specifically, TPR interacts with the TREX2 component GANP, which is required for the export of both intronless and spliced mRNAs (14, 17, 35). Additionally, TPR inhibition by antibody injections, TPR-depletion by RNAi, and overexpression of TPR and TPR fragments, resulted in the nuclear accumulation of poly(A)-RNA – suggesting a role in nuclear mRNA export (11, 36, 37). Moreover, the yeast homologues of TPR, Mlp1/2, interact with the TREX component protein Aly and the poly(A)-binding protein Nab2 (38–41), which has been implicated in mRNA export in yeast. This interaction may promote the docking of mRNP complexes with the nucleoplasmic face of the yeast nuclear pore (42, 43). TREX-Mlp1/2 interactions and mRNP docking could facilitate the exchange of proteins in the mRNP to prepare it for nuclear export (44). Whether TPR associates with TREX components or regulates mRNP remodelling in mammalian cells remains unclear. Since a sizeable fraction of NPC-associated components exist freely in the nucleoplasm, where they are bound to chromatin and regulate gene expression (45–48), it is also possible that soluble TPR may have additional roles in regulating mRNPs even before they reach the nuclear pore.

TPR has also been reported to be required for the nuclear retention of mRNAs with retained introns. In particular, it was seen that TPR-depletion result in an increase of the expression of proteins derived from late HIV transcripts that contain unspliced introns (49, 50). In yeast, deletion of the TPR homologues, Mlp1/2, also led to increased protein expression of an intron-containing reporter mRNA, suggesting again that TPR is required for nuclear retention of such transcripts (51–54). Together this data suggested that TPR may be involved in recognizing and retaining mRNAs that are improperly processed in that they still contain introns. Importantly, these studies have examined protein expression from reporters and never directly observed whether the cytoplasmic/nuclear distribution of these mRNAs was altered. It is likely that mRNAs with retained introns are targeted for nuclear retention by virtue of the fact that they contain intact 5’ splice site (5’SS) motifs (26, 55). Indeed, we previously demonstrated that reporter mRNAs that retain 5’SS motifs in the mature mRNA are nuclear retained and accumulate in nuclear speckles (26). Whether TPR regulates the nuclear retention of these reporters has yet to be directly tested.

Here we show that the nuclear basket protein TPR does not promote the nuclear retention of 5’SS motif containing RNAs, but instead is required for the export of mRNAs and lncRNAs that are generated from transcripts that contain few to no introns. By increasing the number of introns in these pre-mRNAs, their export becomes increasingly independent of TPR. In TPR-depleted cells, these mRNAs accumulate in nuclear speckles and are targeted for decay even though they have recruited Nxf1. Our results indicate that TPR acts at a late step in the mRNA export pathway and may facilitate the passage of certain mRNAs out of nuclear speckles and through the nuclear pore.

## Material and methods

### Plasmids constructs, primers and antibodies

All *ftz* and *βG* reporter construct in pcDNA3.0 were described as previously (24–26, 29). To generate the *USP14-3’UTR* containing plasmids, we amplified a portion of the *USP14* 3’UTR from U2OS genomic DNA using the forward primer: 5’-CTCGAGATTA AGTTTGATGA TGACAAAGTC A-3’ and reverse primer: 5’-TCTAGACAGT CAGAAAATCT AAATTCTAAT TGTT-3’. The ∼ 2.1 kb PCR reaction product was digested using XhoI and XbaI and ligated into the pCDNA3 *ftz* reporter construct that was linearized using the same two restriction enzymes.

Antibodies used in this study are mouse monoclonal TPR-specific antibody (Abcam, ab58344), Nxf1 (rabbit, 53H8, Santa Cruz, sc32319), UAP56 (rat monoclonal (56)), mAb414 (mouse, Abcam, ab24609), Aly (rabbit, Sigma, SAB2702076), tubulin (rat, YL1/2, Invitrogen, MA1-80017), Aly (rabbit polyclonal (2)) for the ER marker, Trap-α (rabbit polyclonal (57)), histone H3 (rabbit polyclonal, Santa Cruz, sc-10809) and histone H2A.Z (rabbit, Millipore, 07-594). All antibodies were diluted 1:1000 for western blotting and 1:250 for immunofluorescence microscopy.

### Cell culture, DNA transfection experiments and Lentiviral delivered shRNA protein depletion

U2OS and HEK293T cells were grown in DMEM media (Wisent) supplemented with 10% fetal bovine serum (FBS) (Wisent) and 5% penicillin/streptomycin (Wisent). DNA transfection experiments were performed as previously described (26).

The lentiviral delivered shRNA protein depletion is as previously described (24, 25, 58). Briefly, HEK293T were plated at 50% confluency on 60 mm dishes and transiently transfected with the gene specific shRNA pLKO.1 plasmid (Sigma), packaging plasmid (Δ8.9) and envelope (VSVG) vectors using Lipo 293T DNA *in vitro* transfection reagent (SignaGen Laboratories) according to the manufacture’s protocol. 48 hours post-transfection, viruses were harvested from the media and added to U2OS cells pre-treated with 8 μg/ml hexadimethrine bromide. Cells were selected with 2 μg/ml puromycin media for at least 4 to 6 days. Western blotting was used to determine the efficiency of TPR depletion. The shRNAs constructs (Sigma) used in this study are as follows: TPR A shRNA, TRCN0000060066 (5’-CCGGGCGATC TGAAACAGAA ACCAACTCGA GTTGGTTTCT GTTTCAGATC GCTTTTTG-3’) and TPR D shRNA, TRCN0000060067 (5’-CCGGCGTAGG TACAAGACTC AATATCTCGA GATATTGAGT CTTGTACCTA CGTTTTTG-3’).

### Microinjection, FISH staining, immunostaining and nuclear speckle Pearson correlation and enrichment quantifications

The microinjection, FISH and immunofluorescence staining and imaging were performed as previously described (24–26, 55, 58, 59). The Pearson correlation quantification was performed as previously described (24). Note that this quantification method has been validated with extensive controls – see (24) Figure 1 D-G. For each experiment, the 10 brightest SC35-positive speckles per cell were analyzed and the totals from 15 to 20 cells were averaged. This analysis was repeated three times and the averages and standard errors between experiments were determined and graphed. For the nuclear speckle enrichment. The nuclear speckle enrichment of *ftz* mRNA was performed as previously described (24). In brief, thresholds were drawn using the SC35 immunofluorescence channel and set so that 10% (+/− 0.5%) of the nuclear area was selected per cell. The fluorescence intensity of RNA in the selected area, in the nucleus and in the entire cell were calculated.

**Figure 1.**
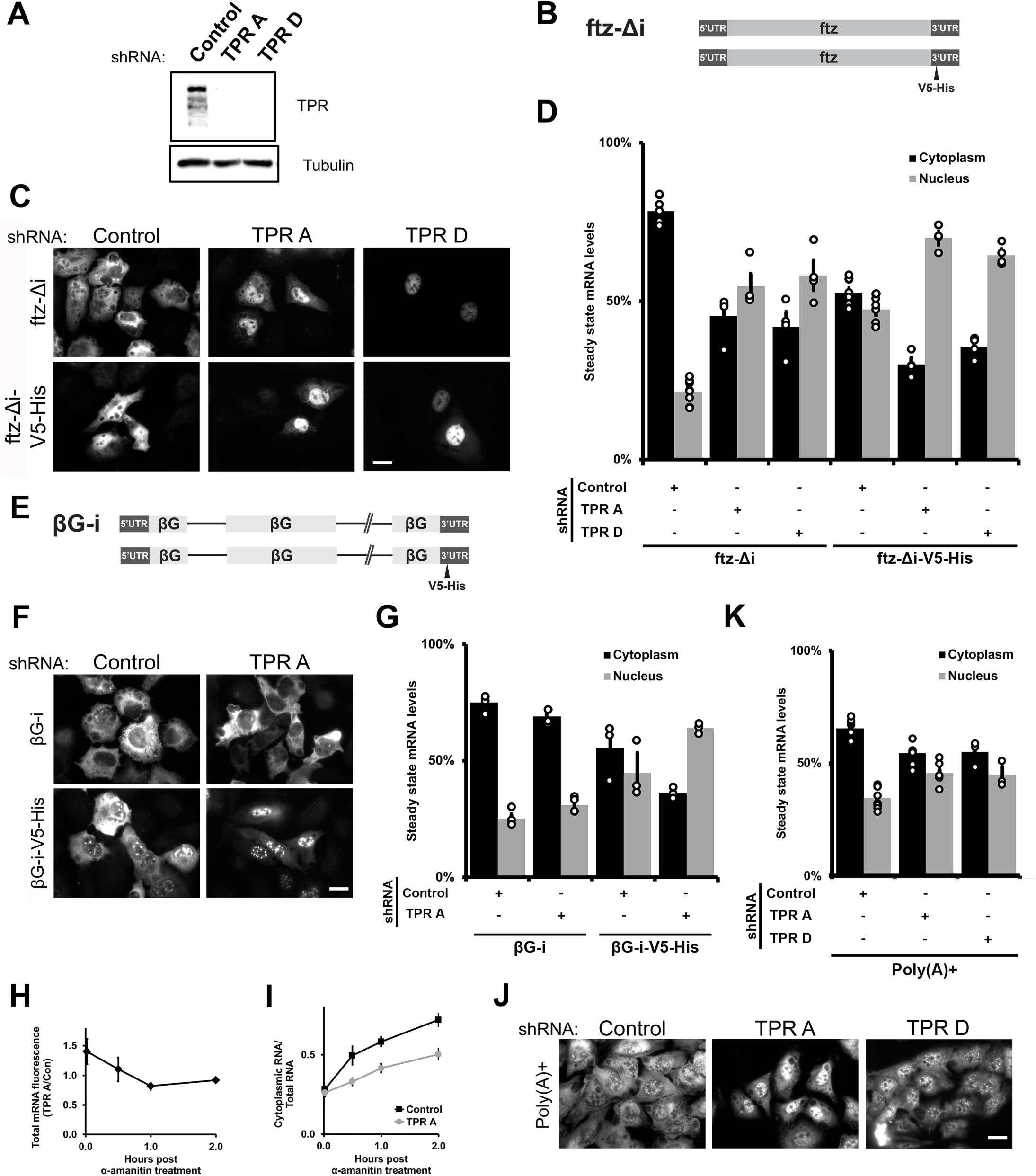
TPR is required for the cytoplasmic accumulation of certain reporter mRNAs but not the nuclear retention of 5’SS motif containing mRNAs. A) U2OS cells were treated with two different lentiviral delivered shRNAs against TPR (“TPR A” and “TPR D”) or scrambled control. Lysates were collected after 96 hrs, separated by SDS-PAGE and immunoprobed for TPR or tubulin. B) Schematic of the intronless *ftz* reporter construct used in this study, with and without the *V5-His* element in the 3’ UTR. The *V5-His* element contains the 5’SS motif which promotes nuclear retention. C-D) Control- or TPR-depleted cells were transfected with the intronless *ftz* reporter plasmid (+/- *V5-His*). 18-24 hours later the cells were fixed and the mRNA was visualized by FISH. TPR depletion caused nuclear accumulation of the *ftz-Δi* mRNA, irrespective of the 5’SS motif. Representative images are shown in (C), scale bar = 10 µm, and quantification is shown in (D) with each bar representing the average and standard error of four independent experiments for either TPR A or TPR D-depleted cells and eight for the control-depleted cells, each experiment consisting of at least 60 cells. E) Schematic of the intron containing *βG-i* reporter mRNA, with and without the *V5-His* element. F-G) Similar to (C-D), except that *βG-i* reporter plasmid was used. TPR depletion did not affect the mRNA distribution of the *βG-i* reporter mRNA but increased the nuclear accumulation of *βG-i-V5-His* mRNA. Scale bar = 10 µm. Representative images are shown in (F), and quantification is shown in (G) with each bar representing the average and standard error of three independent experiments, each experiment consisting of at least 60 cells. H-I) Plasmids containing *ftz-Δi* were microinjected into the nuclei of Control- or TPR-depleted U2OS cells. After allowing mRNA synthesis for 20 min, cells were treated with α-amanitin and mRNA levels were monitored over time by FISH. (H) To determine whether TPR-depletion promotes the degradation of newly synthesized mRNAs, the ratio of *ftz-Δi* signal in the control and TPR-depleted cells were plotted over time. Each point is the average and standard error of at least five independent experiments, each of which consist of 30 to 45 cells. Note that the relative level of *ftz-Δi* in TPR-depleted cells decreases over the first hour until about 60% of the mRNA remains, after which point the ratio is stable. J-K) TPR depletion causes nuclear accumulation of poly(A)+ RNAs, as visualized by oligo-dT FISH staining, scale bar represents 10 µm. Representative images are shown in (J), and quantification is shown in (K) with each bar representing the average and standard error of at least eight independent experiments for Control- or “TPR A” depleted cells and three independent experiments for “TPR D” depleted cells, each experiment consisting of at least 30 cells.

### Stellaris smFISH experiments

Stellaris smFISH experiments were performed as previously described (60) and following the manufacturer’s protocol with a few modifications. Cells on coverslips were washed twice with 1X PBS and fixed with 4% paraformaldehyde (Electron Microscopy Sciences). Next, coverslips were methanol treated for 30 minutes in -20 °C and rehydrated twice in 1X PBS, each time washing for 5 minutes. Cells were washed twice in 2X SSC buffer (150 mM NaCl, 15mM NaCitrate, pH 7.1) with 10% formamide. 100μl of hybridization probe (100mg/ml dextran sulfate, 10% formamide, 2X SSC) containing 125 nM Stellaris probes (LGC Biosearch Technologies) was added to the coverslips and incubated for 48 hours. Subsequently, the coverslips were washed three times with 2X SSC buffer with 10% formamide and mounted onto coverslips using DAPI Fluoromount-G stain mounting solution (Southern Biotech). Cells were imaged and quantified as previously described (60).

### RNA-IP experiments

The Nxf1/TAP RNA-IP experiments were performed exactly as previously described (25, 26), except that TPR-depleted cells were transfected in normal DMEM media without puromycin. For the TPR-FLAG IP experiments, the FLAG-TPR-pcDNA3.0 plasmid (Addgene plasmid # 60882) (61) was co-transfected along with other reporter plasmids. Cells were harvested 18 hours post transfection and the RNA-IP experiments was performed using the same conditions as for the Nxf1/TAP RNA-IP. Superscript IV reverse transcriptase (Invitrogen) was used for Nxf1/TAP RNA-IP and Superscript III (Invitrogen) was used for TPR-FLAG experiments. To access the efficiency of the TPR-FLAG IP, samples were separated on an SDS-PAGE gel and transferred onto a blot for immunoblotting.

### RNA Frac-Seq

2 x 150mm dishes of U2OS cells at 60 to 80% confluency cells were depleted of TPR using shRNA “TPR A” or “TPR D” for 5 days. Cells were isolated by centrifugation at 800 *g,* washed with 1X PBS 3 times and 10% of cell volume was reserved as the “total RNA” fraction. The remaining 90% was resuspended in 500 µl φ buffer [150 mM Potassium Acetate, 5 mM Magnesium Acetate, 20 mM HEPES pH 7.4, 1 mM sodium fluoride, 1 mM sodium orthovanadate, 25X protease inhibitor cocktail (Roche), 1:1000 dilution of SUPERase In™ RNase (Invitrogen) and 0.1% diethylpyrocarbonate]. 500 µl of φ buffer with 1% Triton X-100 (Thermoscientific) and 0.2% sodium deoxycholate was gently added to the resuspended cells and incubated on ice for 3 minutes. The mixture was centrifuged at 800 *g* for 5 minutes and the supernatant (the “cytoplasmic/ER fraction”) was removed and respun at 16,100 *g* for 5 minutes to clear any cell debris. The resulting supernatant was transferred to a new 15 ml falcon tube and 2ml of TRIZOL (Life Technologies) was immediately added. The cell pellet (the “nuclear fraction”) was resuspended with 500 µl of φ buffer, and 500 µl of φ buffer with detergents was added as before and spun at 800 *g* for 5 minutes. The supernatant was removed, the cell pellet was washed with 1 ml of φ buffer and was resuspended in 1 ml of φ buffer. Next, 2 ml of TRIZOL was added to the nuclear fraction, 400 µl of chloroform was added and the tube was thoroughly mixed and centrifuged at 12,500 *g* for 15 minutes. The top layer containing the RNA was removed, washed with chloroform and the RNA was next purified and washed according to manufacturer’s instruction for the RNA PureLink column (ThermoFisher Scientific) column. The same RNA extraction procedure was repeated to extract RNA from the “total” fraction. To monitor the purity of the cell lysate fractions, 50 µl of “cytoplasmic/ER” and “nuclear” fractions was denatured with 1X Laemmli Sample Buffer, separated by SDS-PAGE and probed for various compartment specific markers; Aly as the nuclear marker, Trap-α as the ER marker and tubulin as a cytosolic marker.

To prepare the sample for sequencing, 1 µg of DNase I-treated RNA (Ambion DNA-free DNA removal kit (Invitrogen)) was used. To prepare the RNA library, we followed the manufacturer’s protocol for the NEBNext® Ultra™ II Directional RNA Library Prep kit (NEB, E7765S) and poly(A) selected the RNA using the NEBNext Poly(A) mRNA Magnetic Isolation Module (NEB, E7490S). We used the NEBNext Multiplex Oligos for Illumina (Index Primers #1 - #24) (NEB, E7335S and E7500S) as the unique index primers. RNA quality control was performed at all steps using BioAnalyzer and sequenced at the Donnelly Centre Sequencing facility at the University of Toronto using the Illumnia Hi-Seq500 platform.

### RNA Frac-Seq data processing and analysis

A Salmon index (62) was built from Gencode v.19 protein-coding and long non-coding transcript sequences (63). Fastq files corresponding to each biological replicate were pseudo-aligned to the Salmon index (command: quant --validateMappings). Gene abundances were calculated utilizing Tximport (64). Genes with a mean FPKM > 1 were considered for further analysis and genes from the mitochondrial genome were excluded. Gene-level fold changes in the nuclear to cytoplasmic ratio of TPR and control knockdowns were calculated utilizing DESeq2 (65) (with fold change shrinkage) and compared (formula: log_2_(Nuclear/Cytoplasmic)_TPR(A/D)_ - log_2_(Nuclear/Cytoplasmic)_Control_). Gene-level fold changes in nuclear proportions (formula: log_2_(Nuclear/Total)_TPR(A/D)_ - log_2_(Nuclear/Total)_Control_) and cytoplasmic proportions (formula: log_2_(Cytoplasmic/Total)_TPR(A/D)_ - log_2_(Cytoplasmic/Total)_CTRL_) between TPR and control knockdowns were calculated similarly utilizing DESeq2 (with fold change shrinkage) (65). TPR normalized counts for each biological replicate were calculated utilizing DESeq2 (plotCounts) (65).

Gene length, transcript length (exons-only), total number of exons per transcript, fraction intronic sequence (intronic sequence length per gene/gene length) and exon density (total number of exons per transcript/transcript length) of each transcript were calculated from Gencode v.19 annotation (63) for protein-coding and long non-coding RNA transcripts. A gene-level median of each metric was calculated.

For correlation analysis, spearman correlations of the gene-level median of each metric compared to the fold changes in the nuclear to cytoplasmic ratio between TPR and control knockdowns were calculated. For length bin analysis, protein coding or long non-coding RNA encoding genes were divided into 5 bins based upon median transcript length (less than 1000 bp, 1000–1999 bp, 2000-2999 bp, 3000-3999 bp, 4000-4999 bp, greater than 5000 bp). Genes within each length bin were compared to fold changes in the nuclear to cytoplasmic ratio between TPR and control knockdowns. The relationship between exon number and transcript length was assessed utilizing genes with a median transcript length of (2200 – 2300 bp).

For analysis of the total number of exons, multi-exonic genes were defined as genes with a median total number of exons greater than one and single exon genes were defined as genes with a median total number of exons exactly equal to one (unrounded). Multi-exonic and single exon genes were compared to fold changes in the nuclear to cytoplasmic ratio between TPR and control knockdowns. Analysis of genes in particular bins by the total number of exons was performed by rounding the median total number of exons per gene to the nearest integer, followed by comparison to fold changes in the nuclear to cytoplasmic ratio between TPR and control knockdowns. Histone and ribosomal gene annotation were derived from HGNC (66).

## Results

### TPR is not required for the nuclear retention of 5’SS motif containing mRNAs, but is required for the cytoplasmic accumulation of certain reporter mRNAs

Previously, we identified a *cis*-acting RNA element, the 5’SS motif, that inhibits mRNA nuclear export (26). Since TPR has been implicated in the nuclear retention of mRNAs with retained introns, which contain 5’ SS motifs, we examined whether TPR is required to retain reporter mRNAs with this element. We depleted TPR from U2OS cells using two lentiviral delivered shRNA (“TPR A” and “TPR D”; Figure 1A), and transiently transfected the *fushi tarazu* (*ftz*) reporter mini gene that either contains or lacks a “V5-His” region (Figure 1B), which contains a 5’SS motif that inhibits nuclear export (26). In this case we used a version of the *ftz gene* that lacks an intron (*Δi*). We then visualized the reporter mRNA by fluorescence in situ hybridization (FISH) staining. Unexpectedly, we found that TPR-depletion did not inhibit the nuclear retention of the 5’SS motif-containing *ftz-Δi-V5-His* mRNA (Figure 1C-D). Instead, nuclear retention of this mRNA was enhanced (Figure 1C-D). Interestingly, the cytoplasmic accumulation of the *ftz* reporter that lacks a 5’SS motif (*ftz-Δi*) was also inhibited when TPR was depleted. Thus, TPR is required for the nuclear export of the intronless *ftz* reporter mRNA.

We next tested whether TPR-depletion affected the steady state RNA distribution of another reporter gene, *β-globin* (*βG*), with and without the V5-His region. Note that these constructs contain two introns (*βG-i*; Figure 1E). As was seen with *ftz*, TPR depletion did not inhibit, but rather enhanced, the nuclear retention of the V5-His-containing mRNA (*βG-i-V5-His*; Figure 1F-G). Surprisingly, in contrast to the *ftz* reporter, the distribution of *βG-i* was not significantly affected by TPR-depletion (Figure 1F-G).

From these results we conclude that although TPR is not required for the nuclear retention of mRNAs that have 5’SS motifs, it is required for the cytoplasmic accumulation of certain mRNAs.

### TPR depletion influences mRNA stability, and the rate of nuclear export

The cytoplasmic accumulation of mRNAs can be due to a variety of factors. To determine whether this was due to changes in mRNA decay and/or mRNA nuclear export, we monitored the fate of newly synthesized mRNAs. To generate a pulse of transcribed mRNA, we microinjected plasmids that contained the *ftz-Δi* reporter into control and TPR-depleted cells, which allows for the rapid production of a large number of mRNA molecules in a short period of time (59, 67). After allowing expression for 20 min, we added α-amanitin to halt further transcription and then fixed cells at various time points and determined the amount of mRNA in the nucleus and cytoplasm by FISH.

When we compared the amount of FISH signal in the TPR-depleted versus the control-depleted cells we observed that about 40% of the *ftz* mRNA signal was eliminated within the first hour after transcriptional shut-off, but afterwards the ratio remained relatively constant (Figure 1H). This observation is similar to what we observed with reporter mRNAs that contain nuclear retention elements (26). When the nuclear/cytosolic distribution of the mRNA at theses same time points were analyzed, we observed that the rate of mRNA nuclear export was greater in the control versus the TPR-depleted cells (Figure 1I).

From these experiments we conclude that TPR-depletion causes the nuclear retention of certain mRNAs, and that a fraction of these are targeted for mRNA decay.

### TPR is required for the efficient nuclear export of poly(A)-mRNA

Two previous reports showed that TPR depletion lead to nuclear accumulation of poly(A) signal, suggesting that TPR is required for the nuclear export of a significant number of mRNAs (11, 37). In agreement with this, we also found that depletion of TPR increased the nuclear accumulation of poly(A) (Figure 1J-K). Taken together, our results indicate that TPR likely regulates the nuclear export of many mRNAs.

### TPR regulates the nuclear export of a subset of endogenous mRNAs and lncRNAs but is not involved in the nuclear retention of unspliced mRNAs

Next, we determined which RNAs require TPR for their nuclear export on a global scale using RNA Fractionation Sequencing (RNA Frac-Seq; Figure 2A). Briefly, we isolated nuclear and cytoplasmic/ER fractions from control- or TPR-depleted (TPR shRNA “A” or “D”; Figure 2B, S1) cells, then isolated and sequenced poly(A)+ selected RNAs. Each fraction was relatively free of cross contamination, as nuclear markers were well separated from cytosolic and ER-markers (Figure 2C). To control for changes in the transcriptome observed following TPR depletion, we isolated a “total RNA” fraction and used this dataset to normalize against changes observed for both nuclear and cytoplasmic fractions.

**Figure 2.**
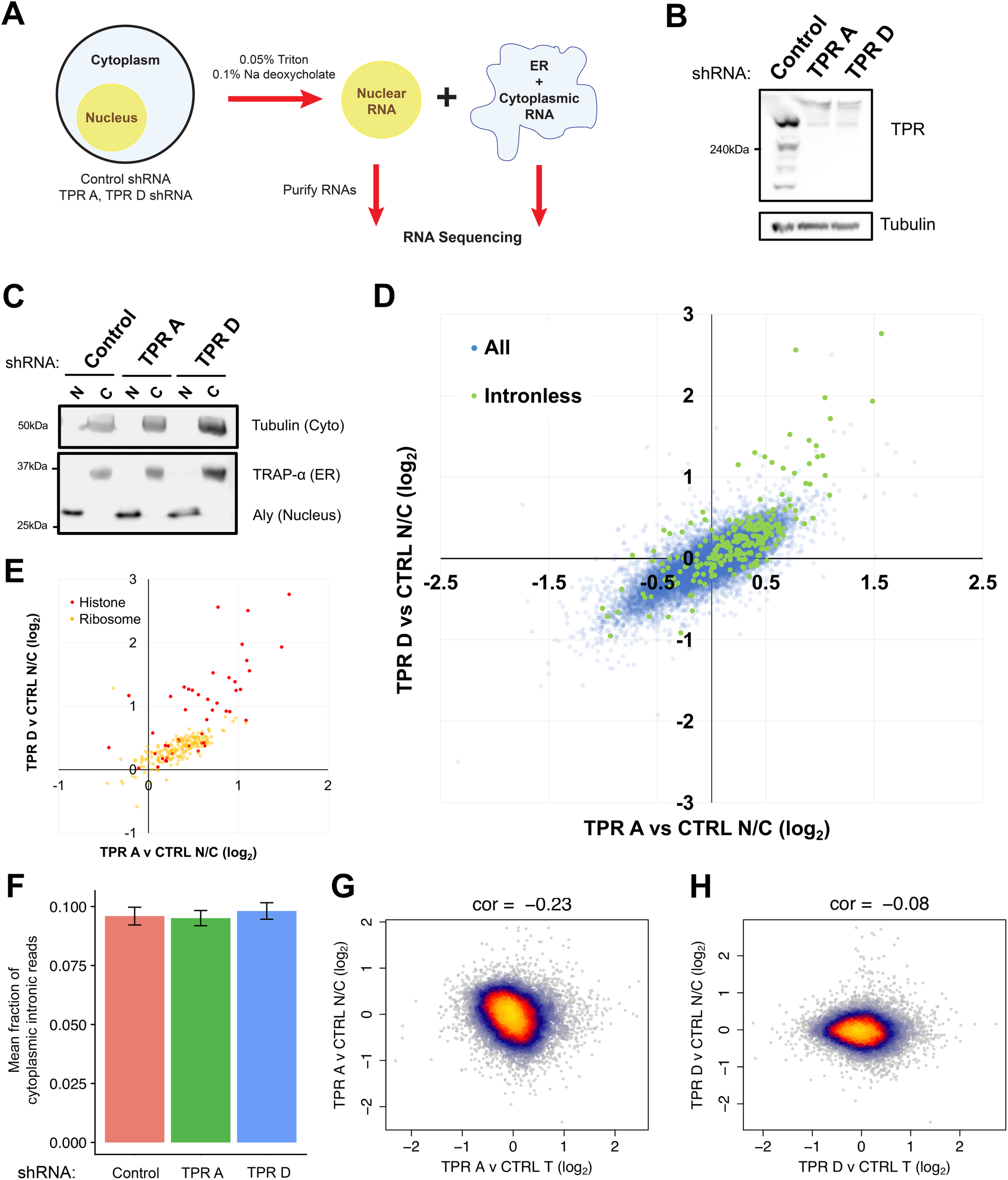
TPR-depletion inhibits the nuclear export of a subset of endogenous mRNAs. A) Workflow for RNA Frac-Seq. Control- or TPR-depleted U2OS cells were fractionated into “nuclear” and “ER/cytoplasmic” fractions (see Material and Methods for more details). RNA from each fraction was purified and sent for RNA sequencing. B) Immunoblot showing TPR-depletion for at least 96 hours post-infection using two different lentiviral delivered shRNAs, “TPR A” or “TPR D”. Tubulin is used as a loading control. C) Immunoblot of nuclear “N” or cytoplasmic “C” fractions, probed with specific nuclear (Aly), ER (TRAP-α) and cytoplasmic (Tubulin) protein markers. D-E) Plot of the Log_2_ fold change of the nuclear/cytoplasmic ratio (TPR v CTRL N/C fold change) between TPR- and control-depleted cells for shRNA TPR A (x-axis) or shRNA TPR D (y-axis) for each mRNA. Positive values indicate an increase in the nuclear/cytoplasmic ratio in the TPR-depleted cells. All mRNA transcripts are shown in (D) with those transcribed from intronless genes (in green) being more nuclear enriched following TPR-depletion compared to those transcribed from other genes (in blue). Particular classes of mRNAs that have high nuclear enrichment upon TPR-depletion are shown in (E), including *histone* (in red) and *ribosomal* (in yellow) mRNAs. F) Fraction of reads that map back to intronic regions of the genome are shown for cytoplasmic fractions from control and TPR-depleted cells. G-H) Plot of the TPR-dependent changes in mRNA level (x-axis) versus TPR-dependent changes in nuclear export (TPR v CTRL N/C fold change) for shRNA TPR A (G) or shRNA TPR D (H) for each mRNA.

To determine which RNAs require TPR for their nuclear export, we calculated the fold change of the RNA isolated from the nuclear and cytosolic compartments. A direct comparison of the nuclear to cytosolic fold change for the TPR-depleted cells against control shows that a subset of RNAs became nuclear enriched upon TPR-depletion (positive fold change value) and this change largely correlated between the two shRNA-treatments (Figure 2D). Upon a cursory inspection, we observed that many mRNAs transcribed from naturally intronless genes became nuclear retained upon TPR-depletion (Figure 2D, green data points). In particular, *histone* mRNAs, which are mostly intronless, were the most affected by TPR-depletion (Figure 2E). Although *histone* mRNAs undergo unique processing events, these mRNAs require TREX for their export (28, 68). Another class of mRNAs that were affected by TPR-depletion were those that encode *ribosomal* proteins (Figure 2E). Unlike histone mRNAs, *ribosomal* mRNAs are generated from genes that have introns, suggesting that it was not only intronless mRNAs that required TPR for their efficient nuclear export.

Next, we analysed the effect of TPR-depletion on the export of mRNAs with retained introns. We observed no significant change in the fraction of cytoplasmic reads that mapped back to intronic regions (Figure 2F), suggesting that TPR depletion did not cause a defect in the bulk nuclear retention of unspliced mRNAs. It however remains possible that small subset of unspliced mRNAs are affected by TPR-depletion.

Recently it was observed that an acute TPR-depletion for 2 hours led to a decrease in the level of certain mRNAs (34). Since we observed that nuclear retained mRNAs were substrates for decay (Figure 1H), we asked whether there was an inverse correlation between mRNA levels and nuclear retention in TPR-depleted cells. However, in our Fraq-Seq data we only saw a weak relationship between changes in mRNA levels and nucleoplasmic/cytoplasmic distribution (Figure 2G-H). Thus, if nuclear retention of an mRNA leads to its decay, this effect is only moderate at most. In addition, compensatory processes may also affect overall mRNA levels in our experiment. For example, we observed that TPR-depleted cells have higher overall levels of histone mRNAs (Figure S2A), and this may explain why overall histone protein levels remain unchanged in TPR-depleted cells (Figure S2B).

In summary our Frac-Seq analysis of TPR-depleted cells indicate that the primary role of TPR is in regulating the nuclear export of a subset of mRNAs. Although it may regulate the nuclear retention of certain mRNAs, TPR-depletion did not promote a detectable increase in unspliced mRNAs in the cytoplasm.

### An analysis of mRNAs that require TPR for their efficient nuclear export

Next, we performed a linear regression analysis to fully explore the features shared by mRNAs that require TPR for their nuclear export (Figure 3A, S3A). Validating our initial observations, we found that the number of exons negatively correlated with TPR-dependent nuclear export (Figure 3A, S3A). In addition, we observed that mRNA and gene length also negatively correlated with TPR-dependent nuclear export. This is not surprising as *ribosomal* and *histone* mRNAs which are both TPR-sensitive (Figure 3B) tend to be short as their encoded proteins are all smaller than 35kDa, whereas the average human protein is about 50kDa. Furthermore, mRNAs generated from intronless genes tend to be shorter than other mRNAs. Moreover, since the primary contributor to gene length is introns, it is also to be expected that intronless and intron-poor genes, tend to be shorter than other genes.

**Figure 3.**
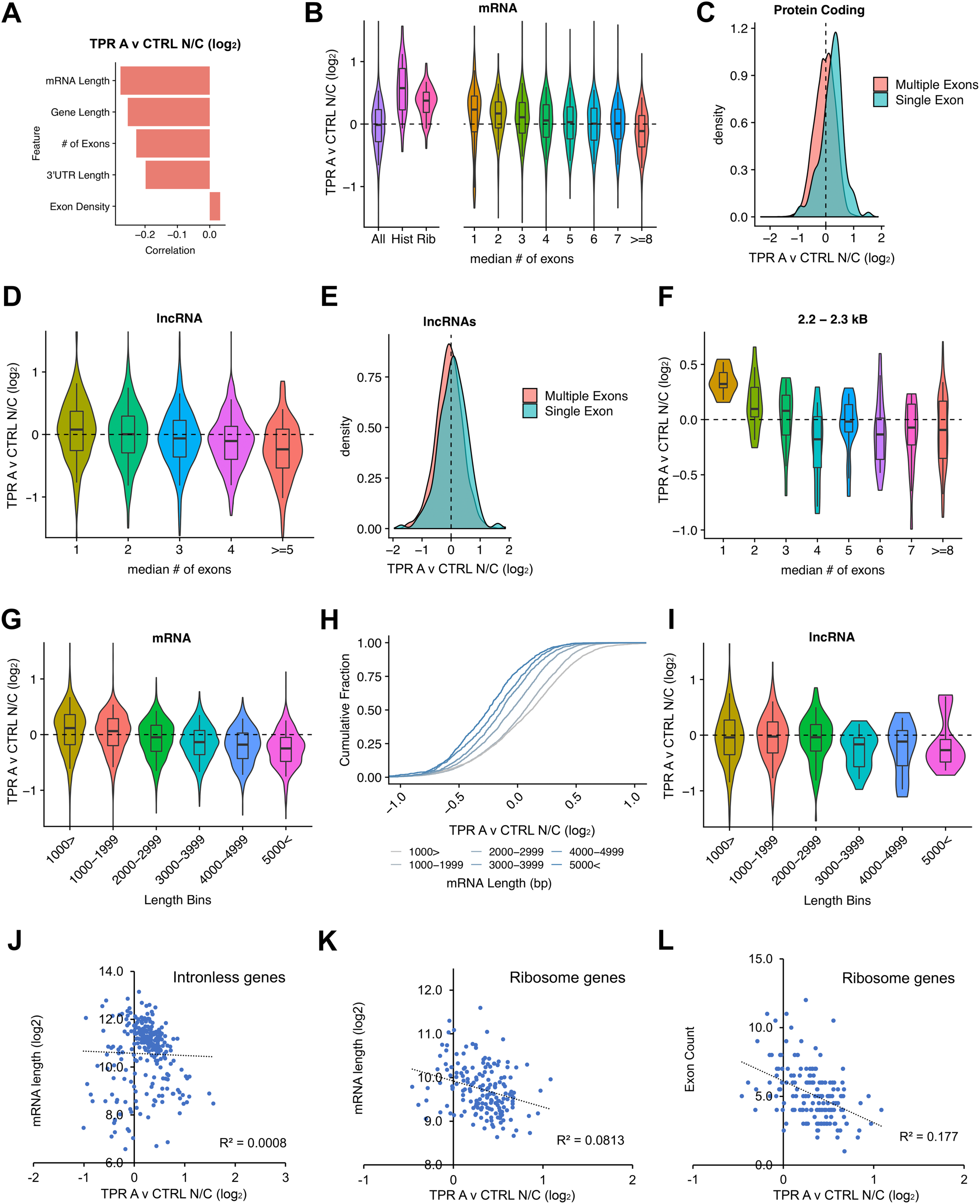
TPR-depletion inhibits the nuclear export of mRNAs and lncRNAs transcribed from intronless and intron-poor genes. A) Regression analysis of TPR-dependent nuclear export (TPR A v CTRL N/C fold change) and various RNA features. Note the inverse correlation between nuclear retention upon TPR-depletion (TPR A v CTRL N/C fold change) and gene length, pre-mRNA length, number of exons and 3’UTR length. B) Violin plots of TPR-dependent nuclear export (TPR A v CTRL N/C fold change) for various classes of mRNAs. Note that mRNAs from *histone* and *ribosomal* genes are more nuclear enriched upon TPR depletion (positive fold change value) compared to all mRNAs. Also note that the nuclear export of mRNAs generated from few exons are more sensitive to TPR-depletion than other mRNAs. C) Density plot showing that mRNAs from intronless genes are more nuclear enriched in TPR-depleted cells compared to those with multiple exons. D) Violin plots of TPR-dependent nuclear export (TPR A v CTRL N/C fold change) for lncRNAs with various numbers of introns. Note that the nuclear export of lncRNAs generated from few exons are more sensitive to TPR-depletion than other lncRNAs. E) Density plot showing that lncRNAs from intronless genes are slightly more nuclear enriched in TPR-depleted cells compared to those with multiple exons. F) Violin plots of TPR-dependent nuclear export (TPR A v CTRL N/C fold change) for mRNAs that are 2.2-2.3 kb in length with various numbers of introns. Again, note the inverse correlation between exon number and nuclear retention upon TPR-depletion. G-I) Violin plots (G, I) and cumulative distribution fraction plot (H) of TPR-dependent nuclear export (TPR A v CTRL N/C fold change) for mRNAs (G-H) and lncRNAs (I) having various lengths. J-I) For mRNAs from intronless (J) or ribosomal protein genes (K-I), the degree of TPR-dependent nuclear export (TPR A v CTRL N/C fold change) was plotted against mRNA length (J-K) or number of exons (I). Note the weak correlation in each case.

In agreement with the idea that splicing (or the lack of splicing) plays a role in TPR-dependent mRNA nuclear export, we found that mRNAs that contained fewer exons were more nuclear-retained upon TPR-depletion (Figure 3B-C, S3B-C). This was also true for lncRNAs (Figure 3D, S3D), however in general there was less of an overall difference between single exon and other lncRNAs in aggregate (Figure 3E, S3E), perhaps because lncRNAs in general have fewer introns.

Since there is a correlation between mRNA length and number of exons, we next wanted to disentangle these two properties. To limit the amount of variation in length, we analyzed mRNAs that are between 2.2-2.3 kb, which is near the average length for the typical human mRNA, and again we observed that the fewer exons an mRNA had the more it was dependent on TPR for its nuclear export (Figure 3F, S3F). Next, we determined whether mRNA length plays a role in TPR-dependent nuclear export. Although we observed an inverse correlation between mRNA length and the inhibition of export after TPR-depletion (Figure 3G-H, S3G-H), we did not see any correlation for lncRNAs that were shorter than 3kb (Figure 3I, S3I). Again, this relationship between length and TPR-dependent export may be due to the fact that shorter mRNAs contain fewer exons. To control for differences in the number of exons, we next looked at the relationship between mRNA length and TPR-dependence for intronless mRNAs. We did not observe a strong correlation between mRNA length and degree of TPR-dependency for export (Figure 3J, S3J), however this may be true for a small subset of intronless mRNAs but that the correlation was obscured because of the many outliers. When we looked at *ribosomal* mRNAs, we did not see any obvious relationship between mRNA length and TPR-dependent export (Figure 3K, S3K), and only a slight inverse correlation between number of exons and TPR-dependent export (Figure 3L, S3L).

Overall, our data suggested that splicing plays a major factor in determining whether the export of a given mRNA requires TPR. However, it remains unclear whether length plays a role in determining TPR-dependency. Importantly, TPR was not only required for the export of mRNAs transcribed from intronless genes, but also from transcripts synthesized from genes with few introns. This agrees with our reporter data, where the nuclear export of the single exon *ftz* mRNA is TPR-sensitive, while the export of the three exon *βG-i* mRNA is TPR-insensitive. In contrast, our reporter data was inconclusive with respect to whether mRNA length plays a role in determining whether an mRNA requires TPR for its nuclear export.

### TPR is required for the efficient export of the *NORAD* lncRNA

Our RNA Frac-Seq results suggest that TPR is required for the nuclear export of mRNAs and lncRNAs that have few exons. To independently assess this, we used single molecule FISH to monitor the distribution of the *NORAD* lncRNA, which is primarily found in the cytoplasm of U2OS cells. *NORAD* is 5378 bp long and has only a single exon, and thus would be expected to be dependent on TPR for its nuclear export. Indeed, we found that upon TPR-depletion, *NORAD* accumulated in the nucleus (Figure 4A-B). In contrast, the distribution of *GAPDH* mRNA, which is 1292 bp long and contains 9 exons, was mostly cytoplasmic in both control and TPR-depleted cells (Figure 4A-B). This data supports the idea that mRNAs and lncRNAs produced from intronless or intron-poor genes require TPR.

**Figure 4.**
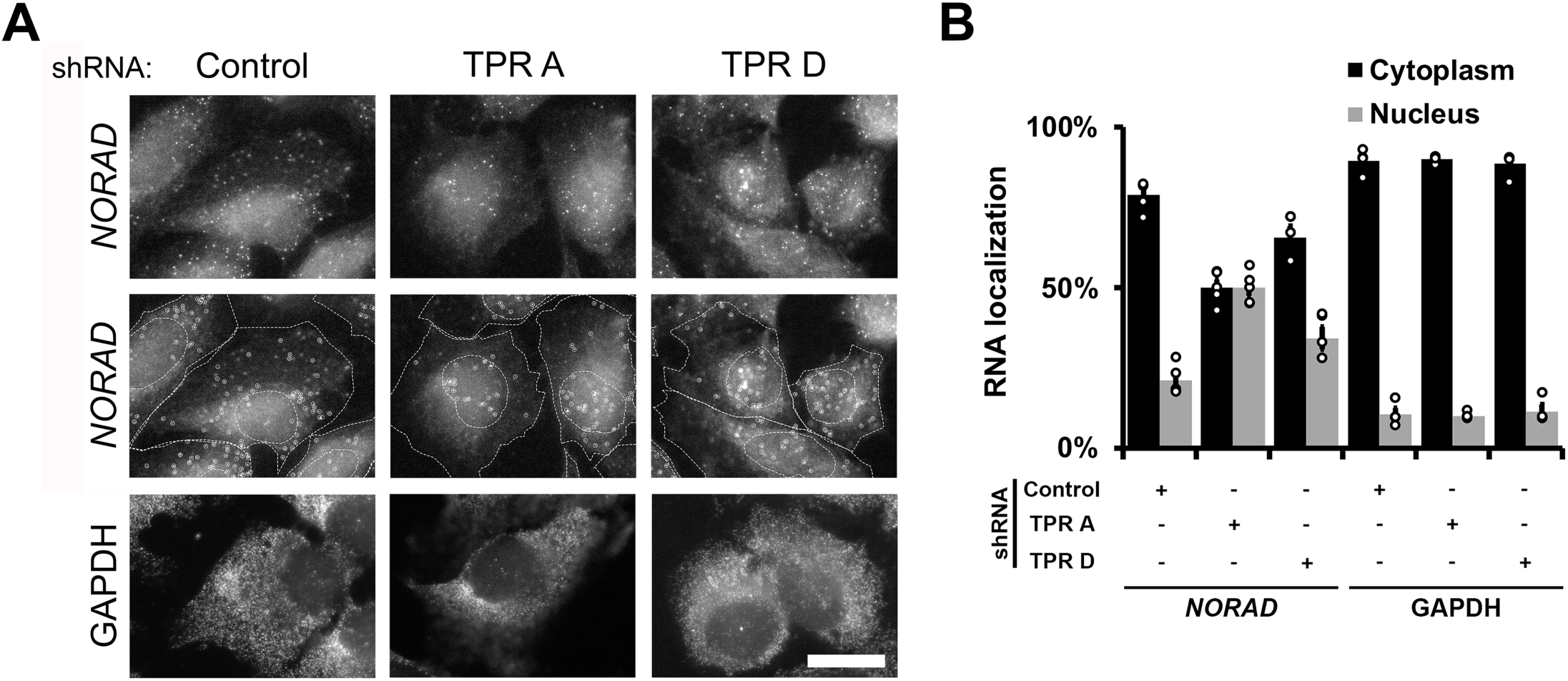
TPR is required for the efficient export of the *NORAD* lncRNA. A-B) U2OS cells were treated with shRNAs against TPR or scrambled control, then fixed and stained for *NORAD* lncRNA or *GAPDH* mRNA by single molecule FISH. Examples are shown in (A) with the top two rows showing the exact same images except that in the second row the cell and nuclear boundaries are delimited by dashed lines and single *NORAD* lncRNAs are circled. The nuclear cytoplasmic distribution of single mRNAs in various conditions is quatified in (B) with each bar representing the average and standard error of at least three independent experiments, each experiment consisting of at least 50 cells.

### Spliced reporter mRNAs are less dependent on TPR for their nuclear export

To further confirm that splicing determines whether an mRNA requires TPR for its nuclear export, we measured the nuclear/cytoplasmic distribution of the *ftz* mRNA synthesized from a reporter that lacked (*ftz-Δi*) or contained (*ftz-i*) an intron (Figure 5A). Indeed, we found that a mRNA produced from *ftz-i* was less nuclear retained in TPR-depleted cells than *ftz-Δi*, suggesting that in this case the presence of one intron partially relieved the need for TPR for efficient export (Figure 5B-C). Interestingly, this mRNA was partially nuclear retained in TPR-depleted cells when compared to control cells. We next wondered whether the presence of two introns would further relieve the need for TPR. Indeed, as we had seen previously, TPR-depletion had no effect on the nuclear export of *βG-i* (Figure 5E), which contains two introns. However, when we tested versions of *βG* that contained a single intron (in this case the *ftz* intron at either the first or second intron site; Figure 5D), mRNA from these constructs were partially nuclear retained in TPR-depleted cells when compared to control cells (Figure 5E).

**Figure 5.**
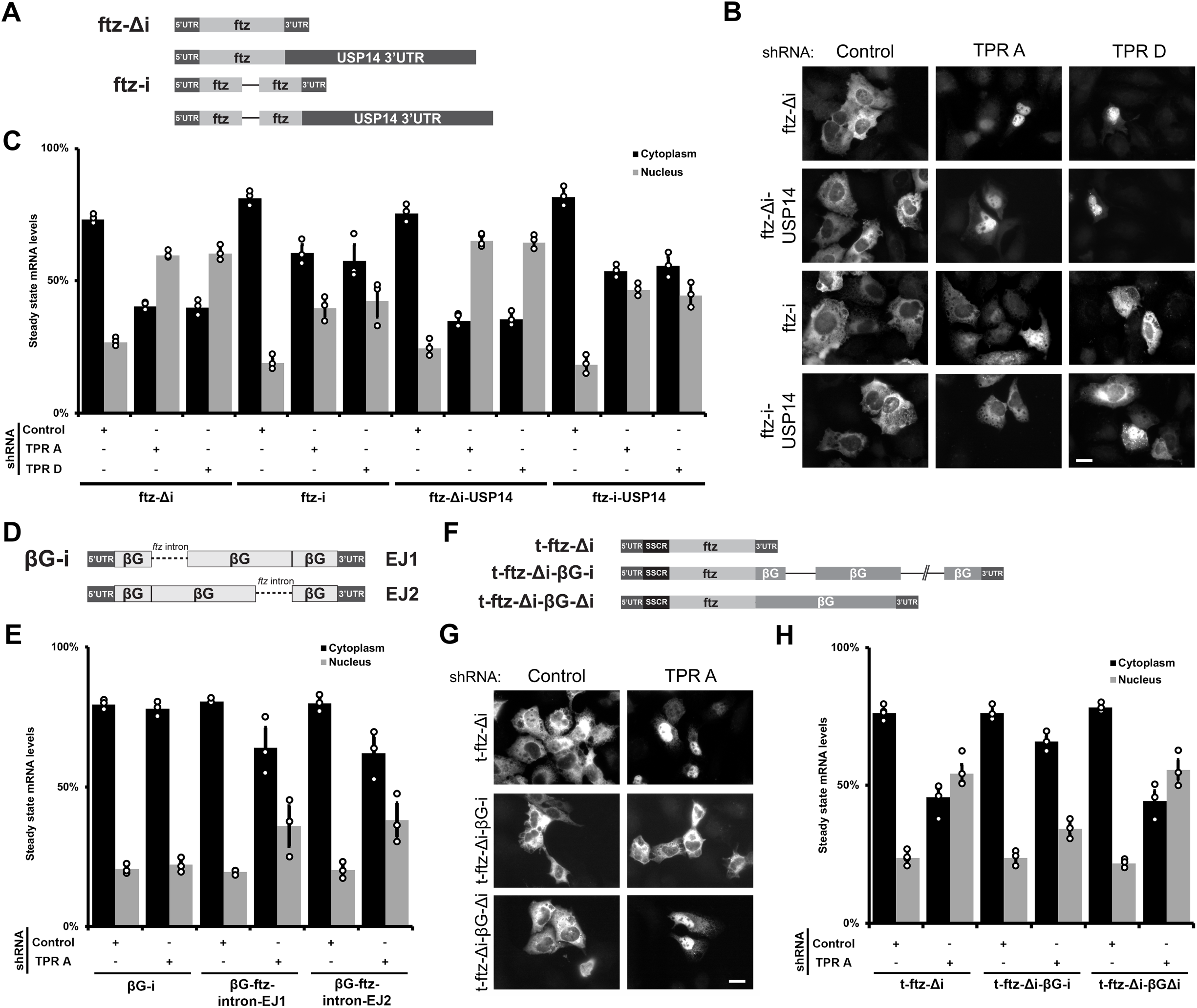
Spliced reporter mRNAs are less dependent on TPR for their nuclear export. A) Schematic of the *ftz* reporter with and without an extended 3’ UTR (+/- *USP14* 3’ UTR), and the *ftz* reporter with and without an intron (+/- *i*). B-C) Control- or TPR-depleted cells were transfected with various *ftz* reporter plasmids. 18-24 hours later the cells were fixed and the mRNA was visualized by FISH. Representative images are shown in (B), quantification is shown in (C). Scale bar = 10 µm. Each bar is the average and standard error of three independent experiments, each experiment consisting of at least 60 cells. Note that TPR-depletion caused nuclear accumulation of the *ftz-Δi* mRNAs, but not the spliced *ftz-i* mRNAs, irrespective of the extended 3’ UTR. D) Schematic of *βG* reporters with the *ftz* intron inserted in either the first exon junction (“EJ 1”) or the second (“EJ 2”). E) Control- or TPR-depleted cells were transfected with various *βG* reporter plasmids. 18-24 hours later the cells were fixed and the mRNA was visualized by FISH and the nuclear and cytoplasmic ratios were quantified. Each bar is the average and standard error of three independent experiments, each experiment consisting of at least 60 cells. F) Schematic of the *t-ftz-βG* fusion reporter mRNAs. G-H) Control- or TPR-depleted cells transfected with the reporters described in (F). 18-24 hours later the cells were fixed and the mRNA was visualized by FISH. Representative images are shown in (G), quantification is shown in (H). Scale bar = 10 µm. Each bar is the average and standard error of three independent experiments, each experiment consisting of at least 60 cells. Note that TPR-depletion caused nuclear accumulation of the *ftz-Δi* mRNAs, but not the spliced *ftz-i* mRNAs, irrespective of the extended 3’ UTR.

In summary, our results indicate that splicing relieves the requirement for TPR to be efficiently exported. Importantly, reporter mRNAs that are have two exons, and are thus singly spliced, are partially sensitive to TPR-depletion for their efficient nuclear export. Thus, it is not just mRNAs produced from intronless genes that require TPR for their export, but also mRNAs produced from genes that have few introns.

### Increasing the length of the *ftz* reporter does not relieve its TPR-requirement for efficient nuclear export

Our RNA Frac-Seq data suggest that mRNA length may determine whether an mRNA requires TPR for its nuclear export, although we could not disentangle this from the fact that short mRNAs also contain few exons. To explicitly test whether the lengthening of an mRNA reduces its requirement for TPR in promoting its efficient nuclear export, we artificially extended the length of the *ftz* reporter mRNA by inserting the *3’UTR* of the USP14 gene, which is 2098 nucleotides long, to generate *ftz-Δi-USP14*, which is approximately 2.7 Kb in length (Figure 5A). Note that the nuclear export of the *USP14* gene is unaffected by TPR depletion. Although this mRNA (*ftz-Δi-USP14*) was efficiently exported in control cells, it was nuclear retained in TPR-depleted cells (Figure 5B-C). As expected, the insertion of an intron (*ftz-i-USP14*), partially restored its export in TPR-depleted cells.

To further test this idea, we next extended the length of the *ftz* mRNA (which is a total of 624 nucleotides long without the poly(A)-tail as determined by cDNA sequencing and 3’cleavage mapping) by inserting a signal sequence coding region (SSCR, 66 nucleotides) to the 5’ end of the ORF (*t-ftz-Δi* Figure 5F) and the *βG* ORF (444 nucleotides) to the 3’ end of the *ftz* ORF (*t-ftz-Δi-βG-Δi*; Figure 5F). We previously demonstrated that these mRNAs are all efficiently exported in a TREX-dependent manner (24, 25). Again, both of these mRNAs required TPR for their efficient nuclear export (Figure 5G-H). Furthermore, the introduction of the two *βG* introns (*t-ftz-Δi-βG-i*; Figure 5F) rescued export of the mRNA in the TPR-depleted cells (Figure 5G-H).

In summary, our experiments with the *ftz* and *βG* mRNAs indicate that the primary determinant of whether a given mRNA requires TPR for its efficient nuclear export is the number of splicing events it undergoes.

### mRNAs that are retained in TPR-depleted cells, accumulate in nuclear speckles

Previously, we and others have observed that in cells depleted of TREX components, mRNAs become retained in nuclear speckles (24, 69), which are small phase-separated regions of the nucleoplasm that are enriched in splicing factors and TREX components (70). Nuclear speckles have been shown to function as sites of post-transcriptional splicing (71, 72). Furthermore, work from our lab and others suggests that many mRNPs are targeted to nuclear speckles, where they either undergo a maturation process to become competent for nuclear export, or are nuclear retained (24, 26, 55, 69, 73, 74). Indeed, it was previously seen that TPR-depletion led to the enrichment of total poly(A) mRNA with nuclear speckles (34, 37). To monitor how newly synthesized mRNAs trafficked through speckles, we microinjected plasmid containing the *ftz-Δi* reporter gene, whose mRNA transits through nuclear speckles (24), and monitored the distribution of this mRNA by FISH and nuclear speckles using antibodies to SC35. In control cells we observed that the newly synthesized *ftz* mRNA quickly entered into speckles, but over time it left these structures as it began to be exported. In contrast, the *ftz* mRNA continued to accumulate into speckles in TPR-depleted cells over the duration of the time course (Figure 6A-C). These observations suggest that TPR may be involved in the release of these mRNAs from nuclear speckles prior to mRNA export. This could be due to direct (i.e., TPR interacting with mRNPs and affecting their maturation) or indirect (e.g., TPR affecting the nuclear/cytoplasmic trafficking of proteins that make up the mRNP) effects.

**Figure 6.**
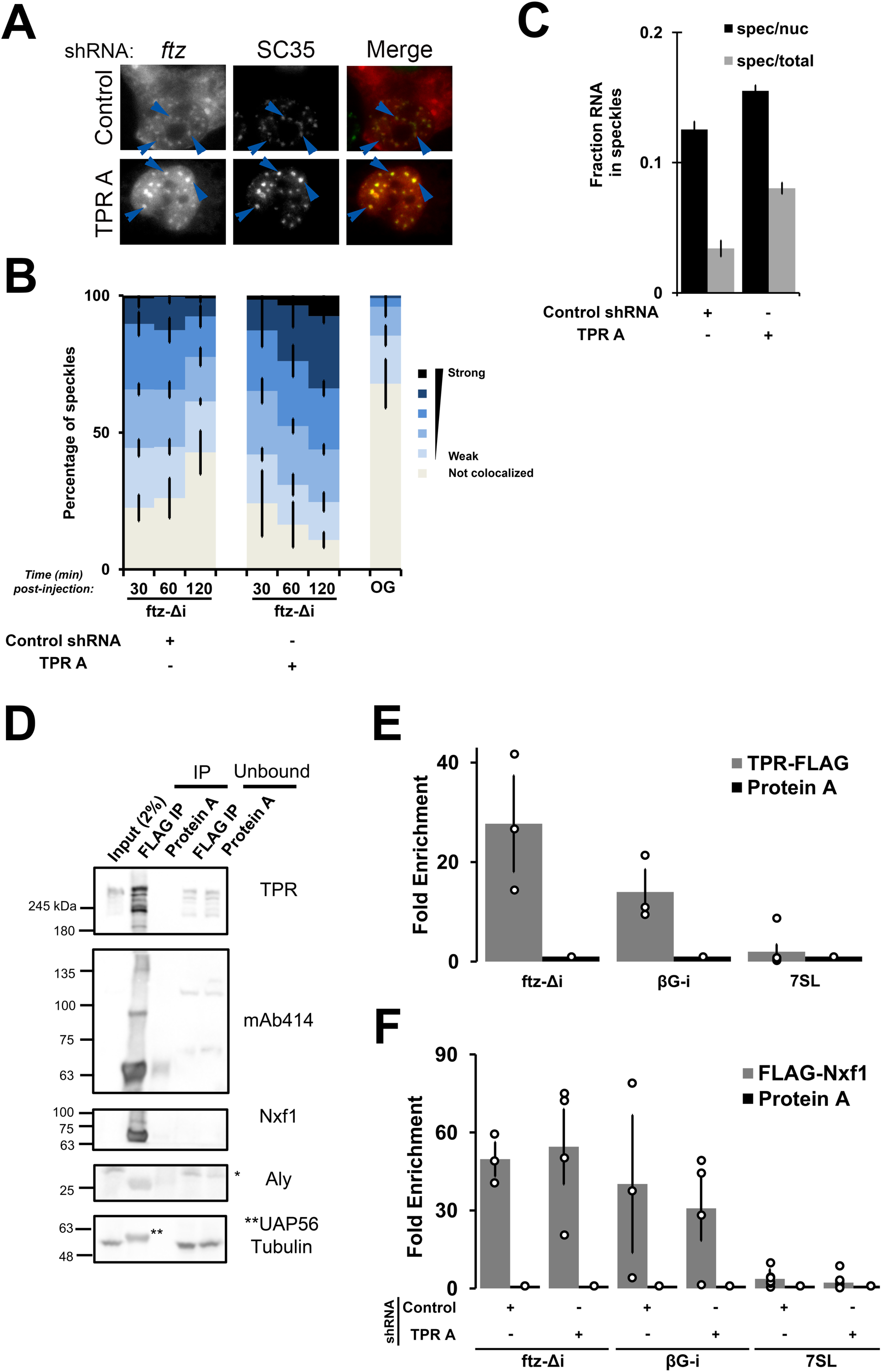
mRNAs that are retained in TPR-depleted cells accumulate in nuclear speckles and are bound to Nxf1. A) Control- or TPR-depleted cells were microinjected with plasmid *ftz-Δi* reporter plasmids. After 2 hours, cells were fixed and stained for *ftz* mRNA by FISH and for the speckle marker SC35 by immunofluorescence. Each row represents a single field of view with blue arrow pointing to examples of *ftz*/SC35 co-localization. B) Control- or TPR-depleted cells were microinjected with plasmid *ftz-Δi* or *ftz-i* reporter plasmids and after various time points were fixed and co-stained for *ftz* mRNA by FISH and for the speckle marker SC35 by immunofluorescence. The degree of *ftz*/SC35 co-localization by Pearson correlation coefficient analysis was quantified as described in (24). As a control, the co-localization of microinjected 70 kD dextran conjugated to Oregon Green (“OG”) with SC35 speckles was also tabulated. Each bar is the average and standard error of three independent experiments, each consisting of 150 to 200 nuclear speckles (see Materials and Methods for more details). C) Control- or TPR-depleted cells were microinjected with plasmid *ftz-Δi* reporter plasmids. After 2 hours, cells were fixed and stained for *ftz* mRNA by FISH and for the speckle marker SC35 by immunofluorescence. The amount of *ftz-Δi* mRNA present in nuclear speckles (defined by the brightest 10% pixels in the nucleus, using SC35 immunofluorescence - described in (24)) as a percentage of either the total nuclear (“spec/nuc”) or total cellular (“spec/total”) mRNA level in cells 1 hr post-microinjection. Each data point represents the average and standard error of the mean of 10–20 cells. D) U2OS cells were transfected with FLAG-TPR for 18 to 24 hours, then lysed and immunoprecipitated with beads conjugated to antibodies against the FLAG epitope or protein A. 2% of the input lysate, Immunoprecipitates (IP), or 10% of the unbound fraction were separated by SDS-PAGE and immunoblotted using the indicated antibodies. Note the presence of unspecific band (*) in the anti-Aly immunoblot. Also note the UAP56 band (depicted as **) in the anti-tubulin and UAP56 immunoblot. E) U2OS cells were co-transfected with TPR-FLAG and either the *ftz-Δi* or *βG-i* reporter plasmids for 18 to 24 hours, then lysed and immunoprecipitated with anti-FLAG M2 beads or protein A beads. The amount of *ftz*, *βG* and the *7SL* ncRNA in the immunoprecipitates were quantified by RT-qPCR. Each bar is the average and standard error of at least three independent experiments. F) Control- or TPR-depleted cells were co-transfected with FLAG-Nxf1 and either the *ftz-Δi* or *βG-i* reporter plasmids for 18 to 24 hours, then lysed and immunoprecipitated with anti-FLAG M2 beads or protein A beads. The amount of *ftz*, *βG* and the *7SL* ncRNA in the immunoprecipitates were quantified by RT-qPCR. Each bar is the average and standard error of at least three independent experiments.

### TPR directly interacts with mRNAs

If TPR directly affects mRNP maturation, then we should be able to detect an interaction. To detect this, we co-transfected our reporter constructs with a FLAG-tagged TPR construct and performed a FLAG immunoprecipitation from the cell lysates. We then detected the presence of the reporter mRNA in the precipitates by RT-qPCR. By immunoblot, we not only detected tagged-TPR in the precipitate, but we also detected other components of the nuclear pore complex using mAb414 antibody (Figure 6D), which recognizes FG-nup proteins found in the nuclear pore (75). The precipitate also contained the mRNA export receptor Nxf1 and the TREX components UAP56 and Aly (Figure 6D). By qRT-PCR we detected both *ftz-Δi* reporter mRNA, which requires TPR for its nuclear export, and *βG-i* reporter mRNA, which is exported independently of TPR (Figure 6E). In contrast, we did not detect the 7SL non-coding RNA (Figure 6E), which utilizes exportin-5 for its nuclear export (76). One interpretation of these results is that TPR binds to all mRNAs during the export process, perhaps mediating mRNP maturation processes. This may be critical for the export of mRNA with few introns, but less important for other mRNAs. This interaction could be occurring at the nuclear pore, where most of the TPR resides, or within the nucleoplasm, where a small pool of chromatin-associated TPR has been detected. When we monitored the distribution of tagged TPR, it was mostly found in the nucleoplasm and not the rim (Figure S4A). This overexpressed TPR did not affect the nuclear export of *ftz-Δi* and *ftz-i* (Figure S4A-B) and did not significantly change the cytoplasmic/nuclear distribution of poly(A)-mRNA (Figure S4A, C).

Following up on the observation that TPR binds to Nxf1 and other mRNA export factors, we tested whether Nxf1 binding to RNA is dependent on TPR. We expressed FLAG-tagged Nxf1 and performed RNA immunoprecipitation experiments in TPR-depleted and control cells. We found that TPR-depletion did not affect Nxf1-binding to either *ftz* or *βG-i* mRNA (Figure 6F). This data suggests that TPR does not help to recruit Nxf1 to the mRNA, but instead acts downstream.

## Discussion

Here we show that the nuclear basket protein TPR is required for the nuclear export of mRNAs and lncRNAs that are produced from either intronless or intron-poor genes. In line with this, we find that the nuclear export of reporter mRNAs becomes more TPR-dependent when they are generated from transcripts with a decreasing number of introns. Furthermore, these mRNAs are associated with Nxf1 suggesting that TPR acts at a very late step in the export pathway. Finally, we observe that these mRNAs become enriched in nuclear speckles, indicating that TPR may influence how mRNAs may traffic through these structures. There has been considerable debate surrounding the role of splicing in nuclear mRNA export (25, 26, 55). What is clear is that intronless mRNAs are exported in a TREX-dependent manner (23, 24, 29, 77), but that they are subject to nuclear retention if they contain certain elements (25). Our new work demonstrates that these mRNAs also have additional requirements for their export, which are less important for mRNAs that are generated by splicing.

Recently, it was found that acute depletion of TPR led to a decrease in the total levels of certain mRNAs (34). This is consistent with the hypothesis that nuclear retained mRNAs are degraded, as is the case with our reporter mRNA (Figure 1H). However, we found that in TPR-depleted cells changes in global RNA levels did not correlate with changes in nuclear/cytoplasmic distribution (Figure 2G-H). Since the depletion was in Aksenova et al., was only for 2 hours, shorter than the average half-life of mRNAs in mammalian cells (5-10 hours (78)), and that TPR did not affect the production of these mRNAs (34), it is possible that the short-term depletion of TPR may regulate mRNA stability, however whether this is associated with nuclear retention needs to be tested. Interestingly, in our experiments the prolonged depletion of TPR led to an increase in the levels of certain nuclear retained mRNAs, such as those encoding histones. This may be a compensation mechanism to ensure that overall histone production remains constant. Another caveat is that prolonged depletion of TPR in our experiments may alter the levels or distribution of factors required for the export of mRNAs transcribed from intron-poor genes and affects their export indirectly.

Our data also indicates that TPR is not required for the nuclear retention of 5’SS motif-containing RNAs. Indeed, we found that if anything, mRNAs with these motifs were even more nuclear retained after TPR-depletion. Previous studies that implicated TPR in the retention of intron-containing mRNAs, investigated viral mRNAs which have additional nuclear retention elements besides the 5’SS motif (55). It thus remains unclear how TPR affects the nuclear retention of these RNAs. Interestingly, one TPR-interacting protein, TARBP2 was found to bind and inhibit the splicing of certain introns that contain TARBP2-binding structural elements(79). The authors proposed that TPR may help to retain these unspliced mRNAs in the nucleus where they are subsequently degraded. When we compared our TPR Frac-Seq dataset, with TARBP2-bound mRNAs (known as the “TARBP2 regulon” (80)) these mRNAs are slightly more cytoplasmic upon TPR depletion (Figure S5A, B), suggesting that TPR may indeed promote their nuclear retention, However, since these mRNAs on average have more exons (S5C, D) and their mRNA is longer in length (S5D) than other mRNAs, it may also be equally likely that these mRNAs are overrepresented in the cytosol of TPR-depleted cells.

The mechanism by which TPR influences export needs to be elucidated. TPR may influence export by interfacing with components of TREX-2. In the absence of TPR, proteins in this complex no longer localizes to the nuclear pore (11, 14, 17, 34). Interestingly, mRNAs that require GANP for their nuclear export tend to be shorter in length and contain few introns (14), mirroring our Frac-Seq analysis of TPR-depleted cells. However, there is some evidence to suggests that TPR and GANP function independently. First, nuclear accumulation of poly(A)+ RNAs was further enhanced in TPR/GANP double-depleted cells when compared to the single depleted cells (14) suggesting that these two factors act on different mRNAs. Second, we previously demonstrated that GANP-depletion inhibited the export of spliced *βG-i* (35), which we now show is exported in TPR-depleted cells (Figures 1F-G, 5E). It remains possible that GANP is required for both TPR-dependent and TPR-independent export. GANP itself has several FG-repeats that may facilitate its partitioning into the central region of the nuclear pore, which is thought to form a phase-separated domain where FG-Nups and nuclear transport receptors partition (81). In light of this, it is possible that TPR may facilitate the interaction of certain mRNPs with GANP, and thus enhance passage through the pore.

In flies, it has been shown that the TPR homolog, Megator, and a second component of the nuclear basket, Nup153, binds to chromatin, forming nucleoporin-associated regions that are dominated by markers of active transcription (47). Similarly, in human cells TPR binds to the promoter region of *HSP104*, and this interaction promotes the association of TPR with the *HSP104* mRNA and is required for its proper nuclear export (82). It is likely that these interactions do not happen in the vicinity of the pore, but within the interior of the nucleoplasm (47), where a substantial fraction of TPR resides (36, 83). Interestingly the ability of TPR to influence how mRNAs traffic through nuclear speckles and to interact with reporter mRNA and components of the TREX complex may suggest that TPR initiates contact with the mRNA before it reaches the nuclear pore.

Finally, it remains unclear why TPR would affect the nuclear export of mRNAs from intronless and intron-poor genes. One factor that may distinguish these is that they tend not to be loaded with exon-junction complexes (EJC), which are recruited to the mRNA during splicing (84). This may indicate that TPR and the EJC act redundantly in promoting nuclear export of mRNAs, and would explain why depletion of EJC components do not generally affect mRNA export (29) despite the fact that they interact with TREX components (84). Alternatively, TPR and the spliceosome may act in a redundant manner to stabilize TREX. In yeast the TPR homologues, MLP1/2, promote the ubiquitination of the yeast Aly, which is required for efficient mRNA export (40, 41). A similar post-translational modification event may be promoted redundantly by TPR and the spliceosome.

## Data Availability

RNA Fraq-Seq data was deposited in GEO database as GSE135537.

## Funding

This work was supported by a grant from the Canadian Institutes of Health Research to AFP (FRN 102725).

## Conflict of Interest

The authors have declared that no conflict of interest exists.

## Acknowledgements

We thank T. Dubric and her staff at the Donnelly Centre Sequencing Facility for performing the RNA sequencing and quality control, B. Cox and S. Pu for their helpful discussions for the analysis of the RNA Frac-Seq dataset and N. Kejiou for providing critical feedback on the manuscript.

---

**Figure S1. (Related to Figure 2) TPR-depletion by shRNA-treatment**

Normalized counts from RNA-Seq showing that TPR was successfully depleted for either total RNA, cytoplasmic or nuclear fractions.

**Figure S2. (Related to Figure 2) TPR-depletion induces the expression of histone mRNAs**

A) Normalized counts from RNA-Seq showing that TPR-depletion induces the expression of *histone* mRNAs.

B) Immunoblots of histone H3 or H2A.Z. from lysates collected from U2OS cells treated with control, TPR A or TPR D shRNA.

**Figure S3. (Related to Figure 3) TPR-depletion inhibits the nuclear export of mRNAs and lncRNAs transcribed from intronless and intron-poor genes**

A-L) Same as in Figure 3, except that the “TPR D” shRNA was used.

**Figure S4. (Related to Figure 6C and D) Overexpression of FLAG-TPR does not influence reporter and poly(A)+ RNA localization**

A) TPR-FLAG was overexpressed in U2OS cells for 18 to 24 hours, then fixed and stained for RNA by FISH and FLAG-TPR by immunofluorescence with antibodies against the FLAG epitope. Note that both poly(A)+ RNA and *ftz* reporter mRNA localization (+/- intron) are not affected by FLAG-TPR overexpression.

B) Quantification of *ftz* reporter RNA and FLAG-TPR protein localization. Each bar is the average and standard error of 45 cells.

C) Quantification of poly(A)+ RNA and FLAG-TPR protein localization. Each bar is the average and standard error of 60 cells.

**Figure S5. TPR-depletion affect the nuclear retention of “TARBP2 regulons”**

A) “TABP2 regulons” are slightly more cytoplasmic in TPR depleted cells (compare orange dots with blue). “TARBP2 regulons” are derived from TARBP2 iCLIP dataset (Fish et al 2019).

B) Average fold change (TPR v CTRL N/C) for “TARBP2 regulons” are cytoplasmic compared to the rest of the dataset (“other”).

C-E) An analysis of “TARBP2 regulon” mRNAs. These have more exons (C) and have longer pre-mRNA (D) than other mRNAs. The average exon length of these mRNAs is similar to that of the entire dataset (E).

