## Supplementary figures and images for "TPR is required for the nuclear export of mRNAs and lncRNAs from intronless and intron-poor genes"

### Supplemental Figures

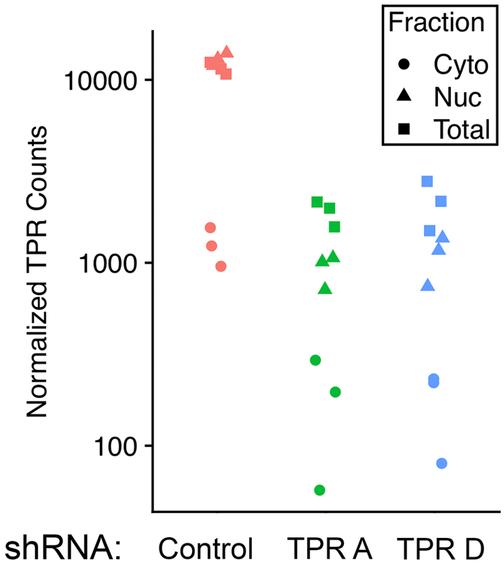

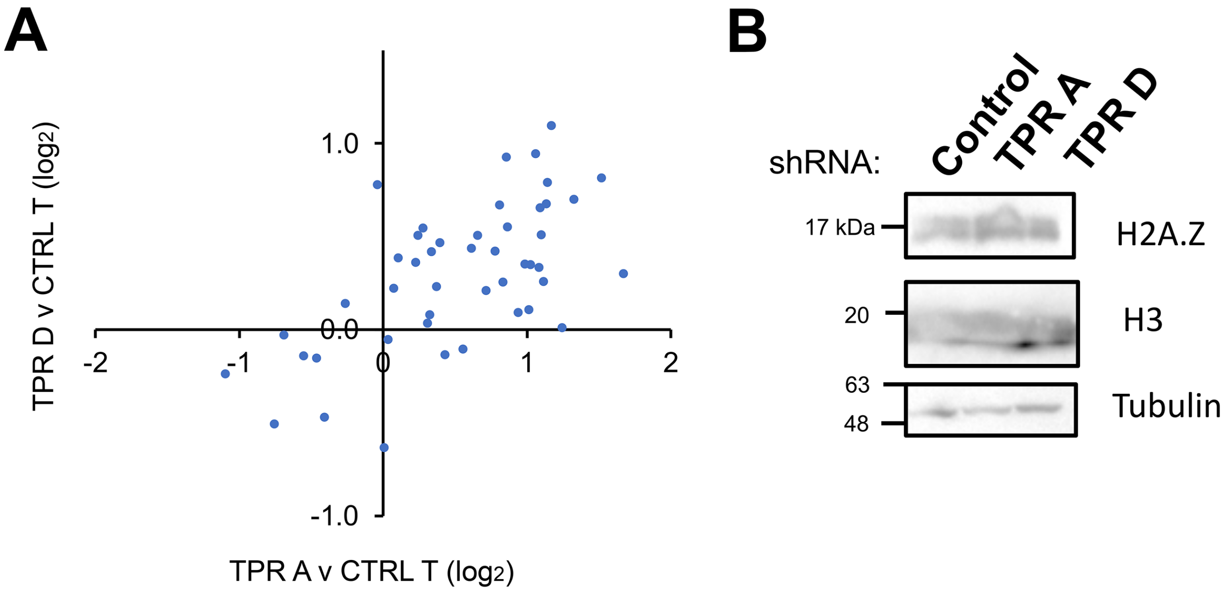

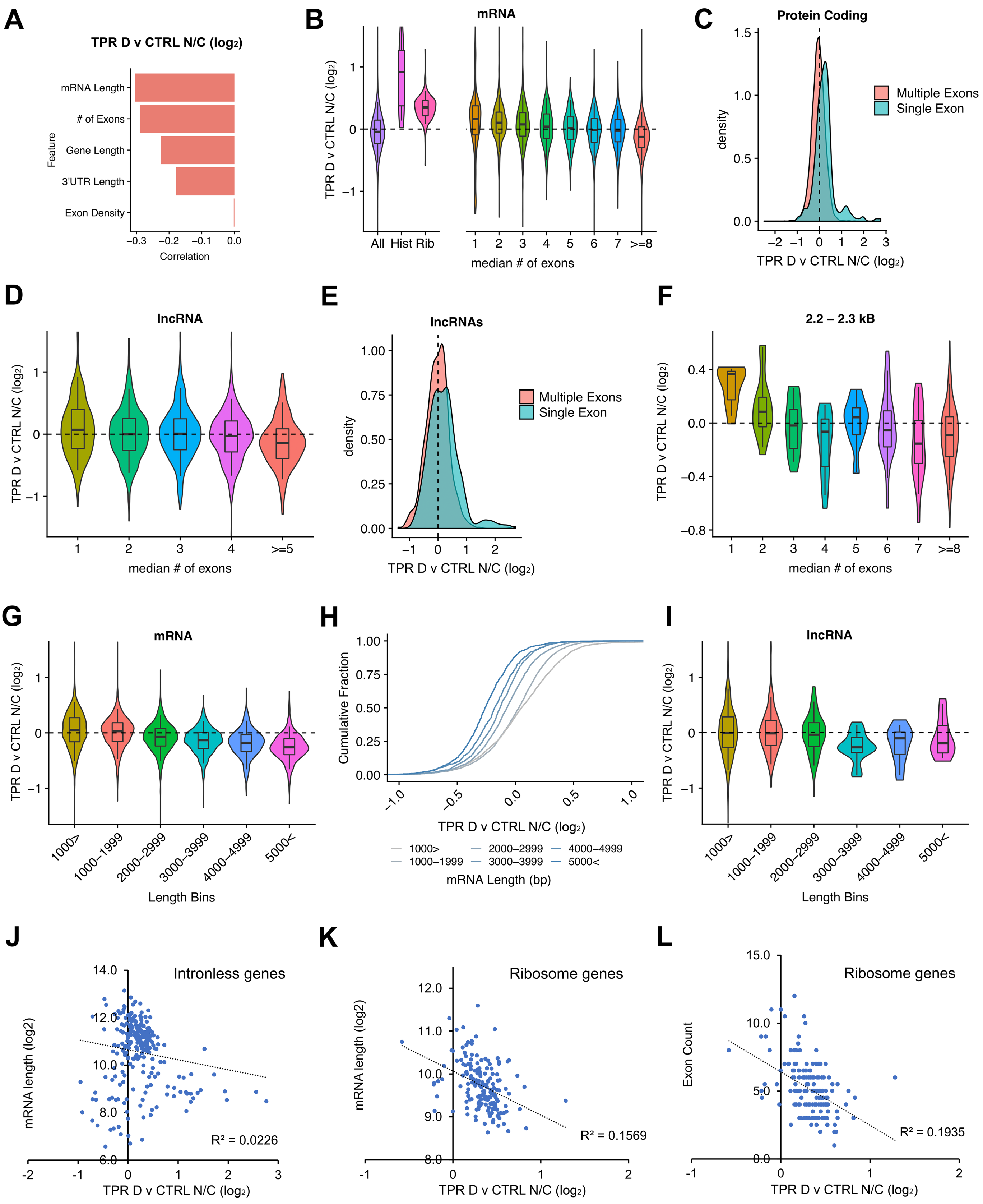

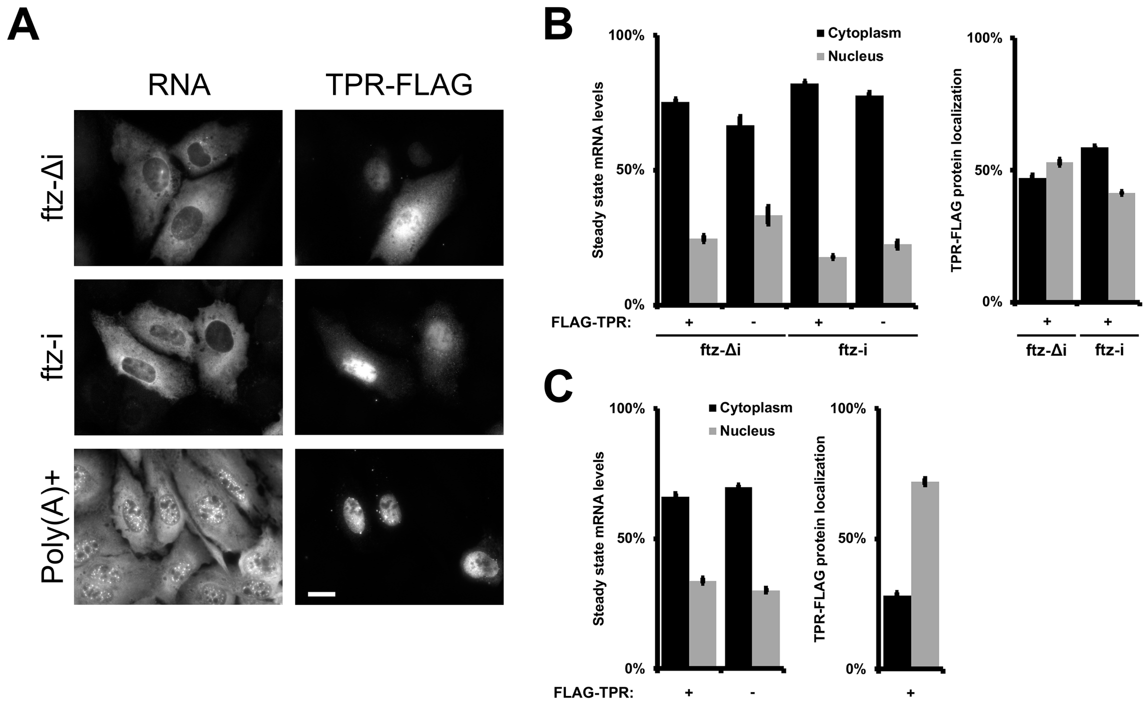

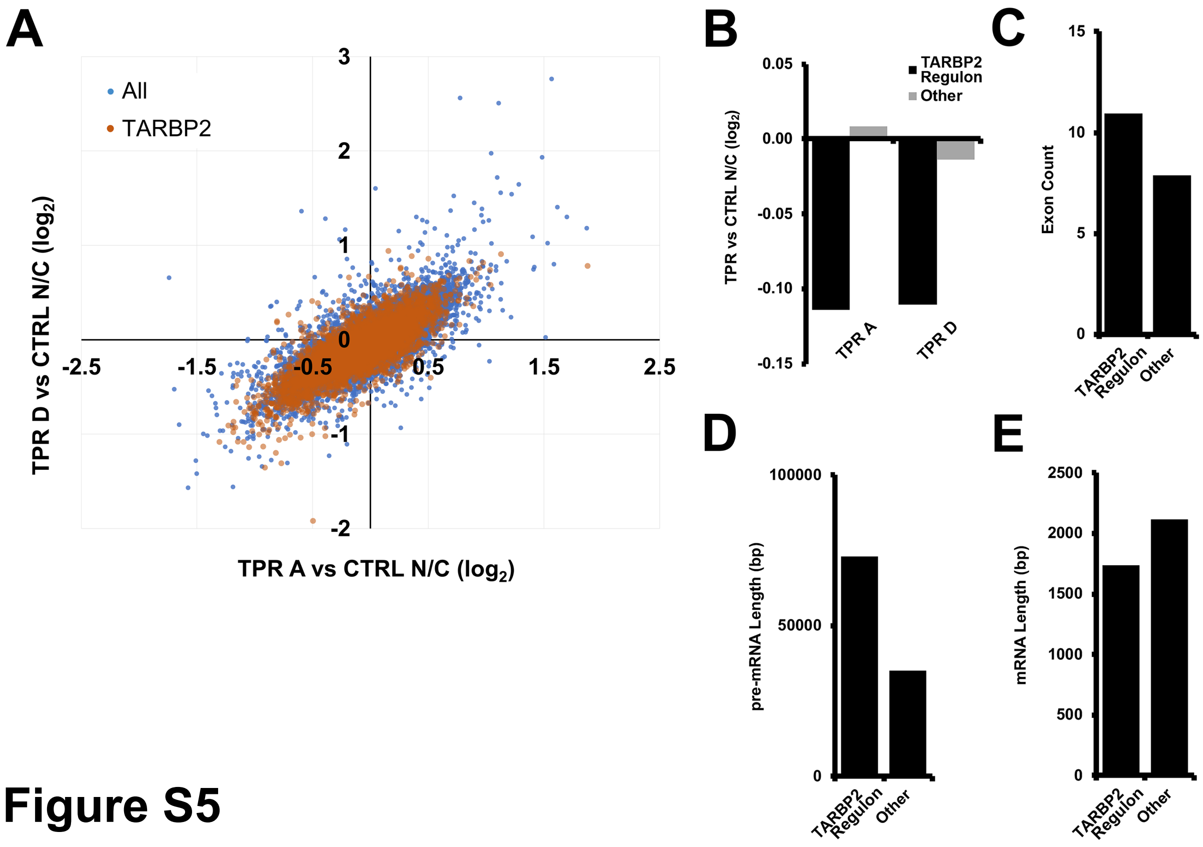
